# Landscape of microenvironment in Randall’s plaque by single-cell sequencing

**DOI:** 10.1101/2021.05.30.444880

**Authors:** Zezhen Liu, Xiaolu Duan, Xinyuan Sun, Jiehui Zhong, Wen Zhong, Bangxian Yu, Zhijian Zhao, Zanlin Mai, Hongxing Liu, Shujue Li, Wenqi Wu, Guohua Zeng

## Abstract

Randall’s plaque is significantly associated with the occurrence of nephrolithiasis. However, the microenvironment of Randall’s plaque is poorly characterized. To investigate the microenvironment of Randall’s plaque, we analyzed single-cell RNA data of 3 Randall’s plaque and 3 normal renal papillae tissue and identified 11 different cell types. We screened differentially expressed genes among all cell types between Randall’s plaque and normal renal papillae. The microenvironment showed two cell types with multiple stone formation-associated transcriptomic programs. Contrary to previous studies, we did not observe macrophage M1/M2 imbalance. Notably, we detected ossification-associated macrophage is enriched in Randall’s plaque and validated GPNMB and ACP5 were potential biomarkers on the ossification-associated macrophage. We also identified an endothelial subset harboring active communication (COL15A1+ PCDH17+ endothelial, DPECs) with other cells. Together with Immunofluorescence, we validated ossification-associated macrophage and DPECs are enriched in Randall’s plaque tissue. Finally, cell-to-cell communication revealed that Loop of Henle, DPECs, and osteoblasts-associated macrophages was the main source of SPP1 signaling. Our work will further the understanding of the microenvironment among Randall’s plaque tissues and provide deep insight into immune modulation.

## Introduction

Kidney stone is a common disease associated with notable morbidity and financial burden affecting all races and ages. The prevalence of symptomatic kidney stone is ranging from 1 to 5% in Asia, 5 to 9% in Europe, and 7 to 13% in North America [1–3]. Kidney stone is also a recurrent disease with a recurrence rate of 50% within 5–10 years and 75% within 20 years [4]. Stone disease is prone to recurrence and there is no effective treatment available, which is related to the fact that the pathogenesis of stones has remained unexplored.

Kidney stones are usually formed as a mixture of crystal aggregates and organic matrix, and they are classified based on major crystalline components [5]. The major components of kidney stones are calcium phosphate (CaP) crystals and calcium oxalate (CaOx) crystals. More than half of all cases with CaOx are referred to as idiopathic CaOx stones. Idiopathic CaOx stones usually develop adhered on Randall’s plaque (RP) existed in kidney papillary surfaces. Randall’s plaque promotes the formation of idiopathic CaOx kidney stones [6], but the detail related genes about stone formation and RP have not been comprehensively identified.

The origin of idiopathic kidney stones has attracted the attention and research by numerous researchers. Microstructure observation found Randall’s plaque originates from the basement membrane of thin loops of Henle in the form of CaP deposits [7]. Taguchi et al. has characterized the genes related to Randall’s plaque formation by microarrays and immunohistology, and found the genes that modulate channels and transporters of sodium/potassium, oxidative stress, vasoconstriction, renal injury, and macrophage [8]. However, the tissue microenvironment of RP remains ambiguous and the exact role and mechanism of RP in CaOx kidney stone formation still lacks in-depth study. The results known so far do not allow for the identification of the type of cell in which the origin of the kidney stone occurs and lack information on the gene expression profile of the specific causative cell. Saeed R. Khan et al. suggested the need to modulate the immune functions with the aim of enhancing polarization of macrophages and hence eliminate crystals, and the potential of blocking plaque growth to suppress it [9]. Recently, single-cell RNA-sequencing (scRNA-seq) has become a comprehensive and central genome-wide method using for determining cellular identities and processes.

In this current study, for uncovering the peculiarities of the Randall’s plaque microenvironment, we performed scRNA-seq on 3 Randall’s plaque from CaOx kidney stone formers and 3 normal renal papillae tissue and identified 11 different cell types. we reveal the differential gene-expression profiling distinct cell types between Randall’s plaque and normal renal papillae. In addition, our work highlights the role of one macrophage subset in Randall’s plaque and activated ECs. These results promote the understanding of Randall’s plaque tissue microenvironment and pathogenesis of some idiopathic CaOx kidney stones.

## Method

### Human specimens

The Randall’s plaque tissues were obtained from patient who underwent ureterscope (URS) at the First Affiliate hospital of Guangzhou Medical University. Before stone removal began, we obtained Randall’s plaque tissue from calyces using BIGopsy Backloading Biopsy Forcepy [10]. Normal renal papillae sample were obtained from three kidney cancer patients who underwent nephrectomy at the First Affiliate hospital of Guangzhou Medical University. Renal papillary tissues were excised from normal adjacent tissues >5 cm distance from the tumor tissue. All subjects were older than 18 years and gave consent to participate. Institutional Review Board approvals were given by the First Affiliate hospital of Guangzhou Medical University (NO. 202091).

### Tissue processing

Single-cell RNA-seq experiment was performed by experimental personnel in the laboratory of NovelBio Co.,Ltd. The tissues were resected stored in MACS Tissue Storage Solution (Miltenyi Biotec). They were processed as described below. First, we washed the tissues in PBS solution and sectioned them on ice into small pieces (approximately 1mm3) followed by enzymatic digestion with collagenase II (Worthington), collagenase IV (Worthington) and DNase I (Worthington) for 40 min at 37°C, with agitation. This was followed by filtration (using 70 μm) and centrifugation at 300g for 5 min. After removal of the supernatant, cell pellet was reconstituted in red blood cell lysis buffer (Miltenyi Biotec) for breakdown of red blood cells. Next, they were washed with 0.04% BSA, and reconstituted in PBS enriched with 0.04% BSA. The suspension was filtered with a 35μm cell strainer. Finally, the cells were used for determination of viability with the Calcein-AM (Thermo Fisher Scientific) and Draq7 (BD Biosciences) after enrichment with MACS dead cell removal kit (Miltenyi Biotec)

### Single-cell RNA sequencing

Single cells were captured with BD Rhapsody system through random distribution of a single-cell suspension across >200,000 microwells by adopting the limited dilution method. Beads with oligonucleotide barcodes were added to saturation so that a bead was paired with a cell in a microwell. After lysing cells in microwells, they were used for hybridization of mRNA molecules to barcoded capture oligos on the beads. We then transferred into a single tube and then reverse-transcribed and ExoI digestion. A cell barcode and unique molecular identifier (UMI) were attached to the 5′ end (that is, the 3′ end of a mRNA transcript) of each cDNA to show its cell of origin. The BD Rhapsody single-cell whole-transcriptome amplification (WTA) workflow including random priming and extension (RPE), RPE amplification PCR and WTA index PCR was employed to prepare transcriptome libraries. Quantification of libraries was done using a High Sensitivity DNA chip (Agilent) on a Bioanalyzer 2200 and the Qubit High Sensitivity DNA assay (Thermo Fisher Scientific). Sequencing was performed by illumina sequencer (Illumina, San Diego, CA) on a 150 bp paired-end run.

### Detection of marker genes

Each cell type (8 cell types) associated with each cluster (25 subclusters) was differentiated from those belonging to other subclusters with the Seurat FindMarkers function. The criteria for marker genes of a cluster were an average expression > 2.5-fold for a given cluster than the average expression in other subclusters from that cell type, and a detectable expression in > 15% of all cells from that subcluster [11]. Moreover, marker genes were those with the highest mean expression in that subcluster, out of all 64 subclusters.

We identified 570 marker genes (Supplementary Table 3) for 25 subclusters (42 stromal cell subclusters), and no marker gene was found for 1 subcluster. The marker genes for many subclusters were collectively analyzed as aggregates, such as for fibroblast, for macrophages and Neutrophil, by combining the marker genes of all subclusters.

### Immunofluorescent staining

Tissue chip, which consists of 7 Randall’s plaque tissues and 5 normal renal papillary tissue, were collected from the First Affiliate hospital of Guangzhou Medical University.

Non-specific binding was suppressed by treat the samples with Immunol staining blocking buffer (Beyotime, cat. no. P0102) for 1 h. The following antibodies were employed for detection: anti-GPNMB (rabbit, 1:50, Proteintech, catalog no. 20338-1-AP), anti-ACP5 (rabbit, 1:50, Proteintech, catalog no. 11594-1-AP) Slides were incubated overnight with antibodies at 4 °C in a moisturized chamber. They were then washed with PBS thrice followed by incubation for 1 h with fluorescence-labelled secondary antibodies (AS001 and AS007, 1:1,000, abclonal) at room temperature in a light-proof moisture chamber. They were then subjected to a three-times wash with PBS, for nuclear staining with DAPI (Beyotime, cat. no.C1002) for 10 min and observed under a fluorescence microscope.

### Sequencing data preprocessing

scRNA-seq data analysis was performed by NovelBio Co.,Ltd. with NovelBrain Cloud Analysis Platform. We applied fastq with default parameter removed low quality reads and filtering the adaptor sequence to achieve the clean data. [12]. The cell barcode whitelist was identified by UMI-tools for Single Cell Transcriptome Analysis. [13]. The UMI-based clean data was mapped to human genome (Ensemble version 91) utilizing STAR mapping with customized parameter from UMI-tools standard pipeline to obtain the UMIs counts of each sample [14]. Cells that contained over 4800 genes or less than 200 were considered discarded or outliers, and Cells in which over 5% of the UMIs were mapped to the mitochondrial genes were discarded. Seurat package (version: 3.1.0, https://satijalab.org/seurat/) was used for single-cell normalization and harmony was used to correct for batch-effects [15]. The top 2000 high variable genes were used to construct PCA and top 15 principals were used for tSNE construction. Graph-based clustering were performed based the top 15 principals to acquire the unsupervised cell cluster. the marker genes were calculated by FindAllMarkers function with Wilcox rank sum test algorithm under following criteria: 1. P value < 0.05; 2. log2(Foldchange) > 0.25; 3. min.pct > 0.15. In order to identify the cell type detailed, re-tSNE analysis, graph-based clustering and marker analysis were performed again on the clusters of same cell type.

### Pseudotime analysis

We applied the (http://cole-trapnell-lab.github.io/monocle-release) Under the default parameters and DDR-Tree settings [16]. we employed the Single-Cell Trajectories analysis utilizing Monocle2 platform to conduct pseudotime analysis. Prior to the analysis, specific genes associated with Seurat clustering were selected from which raw expression counts for the cells were filtered. The 2D tSNE plot was utilized to visualize the trajectories, whereas the plot_pseudotime_heatmap function was employed to create dynamic expression heatmaps.

### Characterization of genes showing differential expression

The FindMarkers (two condition comparison) and FindAllMarkers (multiple condition comparisons) of Seurat package were utilized uncover differentially expressed (marker) genes for each cluster or subtype. Significant Gene markers with differential expression were chosen based on the adjusted P values <0.05, average fold-change >1.5 and percentage of cells with expression > 0.1.

### Cell communication analysis

To enable a systematic analysis of cell–cell communication molecules, we applied cell communication analysis based on cellchat (v1.1.0). database CellChatDB is a manually curated database of literature-supported ligand-receptor interactions [17]. Number of interactions and interaction weights were calculated based on the secreted signaling database and the normalized cell matrix achieved by scran normalization. Visualize cell-cell communication mediated by multiple ligand-receptors or signaling pathways. Systems analysis of cell-cell communication network identify signaling roles of cell groups as well as the major contributing signaling.

### Gene set variation analysis (GSVA)

To evaluate pathway enrichment for different clusters, Gene Set Variation Analysis (GSVA) was predominantly performed using the GSVA R package on the 50 hallmark pathways described in the molecular signature database, exported using the GSEABase package (version 1.36.0) [18]. We also assessed metabolic pathway activities, macrophage pathway activities and associated-ossification pathway activities using a described curated dataset.

### Transcription factor analysis

To assess TF regulation strength, The SCENIC analysis was run as described that passed the filtering, using GRNboost (R package: SCENIC version 1.2.4, which corresponds to RcisTarget 0.99.0 and AUCell 0.99.5; with RcisTarget.hg19. motifDatabases.20k) and the 20-thousand motifs database for RcisTarget [19]. The normalized expression matrix from Seurat were used input of SCENIC.

### Analysis of differential pathway or regulon activities

To quantify differential activities of pathway (GSVA) and regulons (SCENIC) between subsets. We used generalized linear model to contrast the activity scores for each cell. To avoid inflating signals because of interindividual differences, we always included the patient of origin as a categorical variable. Results of these linear models were visualized using bar plots or heatmaps. For the latter, pathways or regulons that did not show significant changesin any of the sets of cells contrasted in one analysis were not visualized (Benjamini–Hochberg-corrected P value > 0.05).

### Gene set enrichment analysis (GSEA)

GSEA analysis was performed to detect which gene set was significantly enriched. Rank-ordered gene lists derived from Pearson correlation coefficients between SPP1 and individual gene expression levels across all Loop of Henle cells were subjected to gene set enrichment analysis[20, 21]. Only gene sets with false discovery rate (FDR) p values <0.05 and nominal p values <0.05 were considered significantly enriched.

### Assessment of osteoblasts activity in macrophage

Osteoblasts-associated genes were obtained from previous study, which included ACP5, CTSK, NFATC1, MAPK1, AKT1, TNFSF11, TRAF6, NFATC1. The mean for osteoblasts-associated genes was adapted to assess osteoblasts activity.

### Statistics

No statistical method was used to predetermine sample sizes. For all experiments, samples from a single patient were processed in parallel, and cells for each sample of one patient were processed for scRNA-seq at the same time, but in separate lanes and vials. Spearman correlation analysis was performed using the R package, and the results were then visualized as scatterplots. Comparisons between two groups were done using two-sided t-test or two-sided Wilcoxon test. P-value were adjusted using Bonferroni correction. Multiple group comparisons were done with One-way analysis of variance (ANOVA).

## Results

### scRNA-seq and cell typing of Randall’s plaque and normal kidney papilla

We obtained 3 plaques from consecutive patients with pure or predominantly calcium oxalate (CaOx) stones who had undergone endoscopic stone removal and 3 normal renal papilla tissues from consecutive patients with kidney cancer who had undergone nephrectomy. Following resection, the tissues were immediately lysed to form single-cell suspensions which were examined with scRNA-seq (see Methods) (**Fig1. a**). Finally, ~0.2 billion unique transcripts were identified from 22947 cells which had more than 100 expressed genes. Among them, 13166 cells (57.4%) were due to Randall’s plaque and 9781 cells (42.6%) from normal kidney papilla. Next, the principal component analysis (PCA) was performed on the genes that were altered across the 52,698 cells (n= 2000 genes) (**Fig1. b**). They were grouped by graph-based clustering based on the important principal elements (n = 20). Finally, cell clusters which were linked to known cell lineages were identified based on markers. Moreover, immune cells (mast cells, macrophages, plasma cells, monocytic, dendritic, T and B cells), fibroblasts, endothelial cells (ECs), epithelial cells and SMC were identified for kidney papilla cells (**Fig1. c, Supplement Table 1**) [22]. The cell type composition of individual Randall’s plaque differs substantially. These differed considerably in transcriptional activity, as we detected on average 8522 transcripts (885 genes) for each Loop of Henle and 4,546 transcripts (2585 genes) for each EC (**Fig1. d**).

### Macrophage tend to be ossification in the microenvironment in Randall’s plaque

Macrophages are linked to interstitial crystals in kidney papillae [23]. To further characterize the macrophages, the level of marker genes was quantified. The frequency of CD14^+^ CD163^+^ CX3CR1^+^ M2 macrophages was considerably higher, whereas the frequency of CD14^+^ CD68^+^ CCR2^+^ M1 macrophages was near absent (**sFig. 1a**). The alternative (M2) type and classic (M1) type are known to affect RP formation, with M2 having an inhibitory effect on inflammation and reducing effect on CaOx crystal deposition [24]. Macrophages were re-analyzed, yielding 5 clusters (**Fig. 2a, b**). Notable heterogeneity exists among macrophage clusters. The CD163, CD86 and CD14, were elevated in all clusters. The C1q protein family genes, APOE and TREM1 are enriched in clusters 0, 2, and 4 [25]. This shows that cluster 0, 2, 4 represented an M2-like macrophage cluster. Additionally, clusters 1, 3 are high in VCAN, FCN1 and S100A protein family genes, which encode calcium-binding proteins (**Fig. 2c**). Activated mononuclear cells secrete proteins with enhance inflammation implying that clusters 1 and 3 may play trigger inflammation in renal papillae [26]. Results from GSVA analysis confirmed that interferon (IFN), complement and inflammatory response pathways were enriched in M2-like macrophage from Randall’s plaque, which are the hallmarks of M2-like macrophages (**Fig. 2d**). SCENIC results confirmed that genes influenced by FOSL2 was upregulated, whereas the EGR1 regulons were decreased in the context of pro-inflammatory macrophage, as similar to a previous report (**Fig. 2e**) [26]. No differences were evident between the RP and NP groups with regards to pro-inflammatory and anti-inflammatory macrophage infiltration. (**sFig. 1b**). We observed a mixture of M1 and M2 signal activation among macrophages, with a strong positive correlation between the two signals (**Fig. 2f**) [27]. Macrophage co-high express M1 and M2 signals in Randall’s plaque. But no significant difference was observed for the M1/M2 signal (**sFig. 1c**). All of this data reveal imbalance of M1 and M2 polarization is not evident in Randall’s plaque compared normal renal papillae. Moreover, the genes associated with ossification changes showed a remarkable expression in Cluster 0, 3, 4 (**Fig. 2g, sFig. 1d**). Specifically, VIM, SPP1, MGP, ACP5 and FN1 associated with ossification in the kidney.

**Fig. 1.**
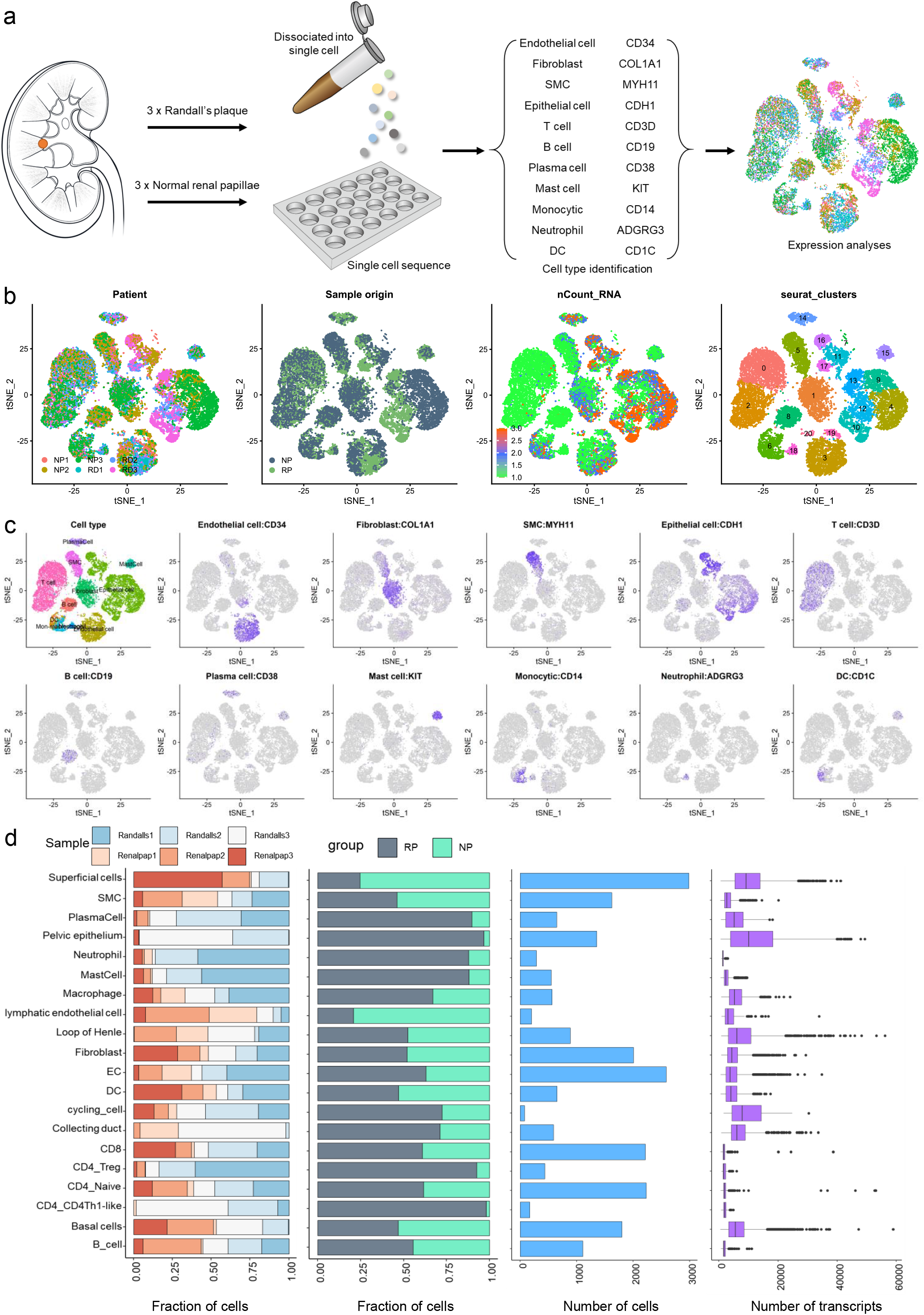
Overview of the 22947 single cells from Randall’s plaque and normal renal papillae samples. a. Scheme of the overall study design. Single-cell RNA sequencing was applied to Randall’s plaque tissue and normal renal papillae.
b. tSNE of the 22947 cells profiled here, with each cell color-coded for (left to right): the corresponding patient, its sample type of origin (Randall’s plaque tissue and normal renal papillae), The number of transcripts (UMIs) detected in that cell, the cell type.
c. Expression of marker genes for the cell types defined above each panel. Three additional marker genes for each cell type are shown in **Supplementary Table 1**.
d. For each of the 20 cell types (left to right): the fraction of cells originating from 6 patients, the fraction of cells originating from each of the 3 Randall’s plaque and normal renal papillae, the number of cells, and the box plots of number of transcripts.

**Fig. 2.**
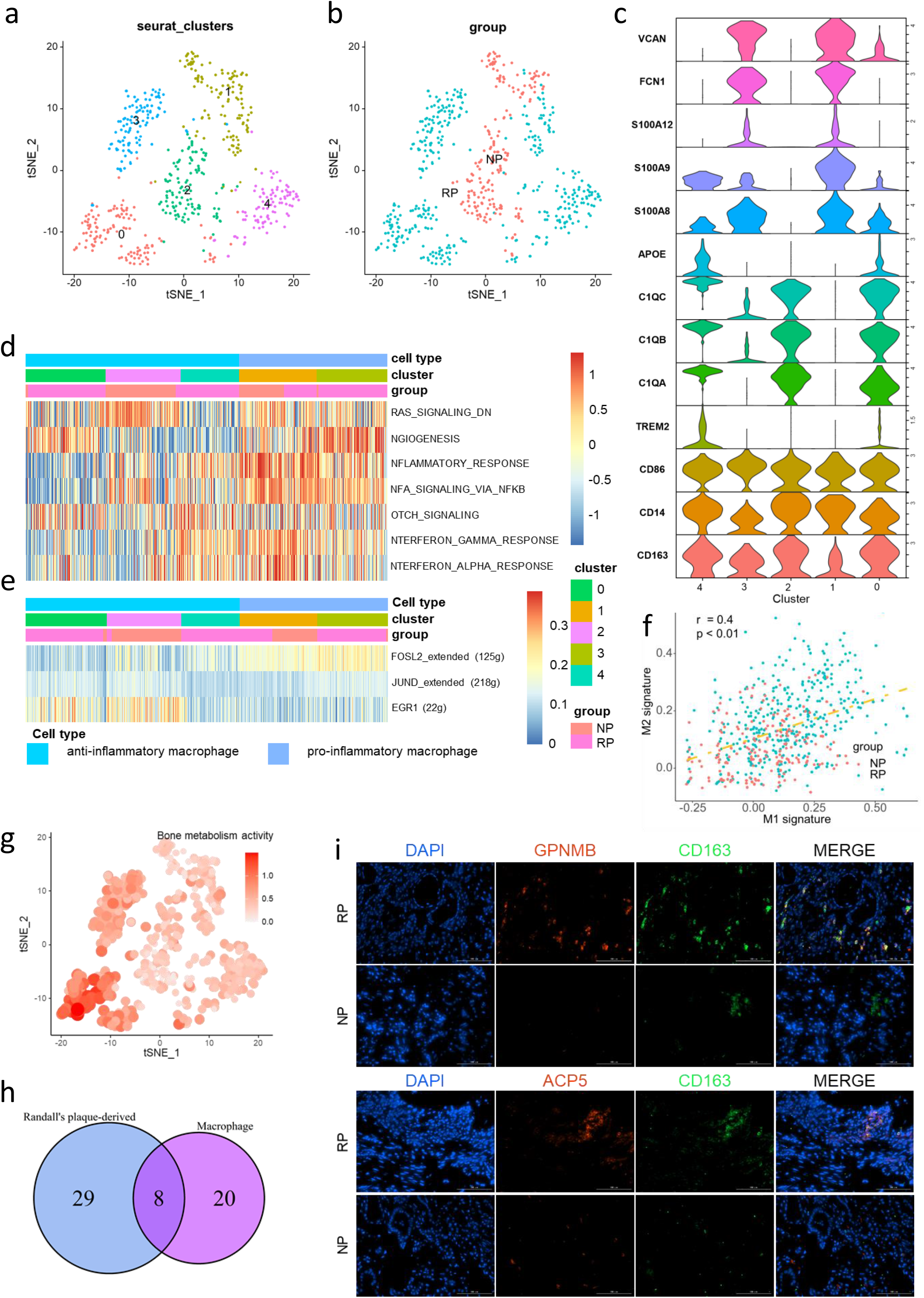
Identifying the characteristics of RP macrophage. a. tSNE plot of macrophage, color coded for 5 clusters of macrophages.
b. tSNE plot of macrophage, color coded for macrophage from Randall’s plaque and normal renal papillae.
c. pro-inflammatory and inti-inflammatory specific markers of macrophage.
d. Differences in 50 hallmark pathway activities scored with GSVA between RP and NP. Shown are t values calculated by a linear model.
e. Heatmap of AUC scores of selected regulons altered in cancer cells. AUC scores were measured by SCENIC per cell.
f. Correlation between the M1 and M2 signatures, where each dot represents a cell. Two-sided P values are calculated for Spearman’s rank correlation.
g. The distribution of osteoblasts activity in macrophage. The color scale represents mean of osteoblasts associated-genes.
h. The Venn diagrams show the number of differentially and commonly markers identified between Randall’s plaque-derived macrophage and all macrophage.
i. Immunofluorescence staining of Randall’s plaque and normal renal tissue chips. GPNMB and ACP5 were co-stained with CD163 in Randall’s plaque, respectively. Scale bar represents 50 mm.

To identify Randall’s plaque-specific marker genes for macrophage, elevated genes in Randall’s plaque-derived macrophage were overlapped with those primarily expressed in macrophage. This resulted in 8 genes, with ACP5 and GPNMB displaying the best specificity (**Fig. 2h, sFig. 2a**). ACP5 is involved in osteoblast regulation and macrophage function, as well as reflecting bone resorption and osteoclast activity [28]. GPNMB has been linked to osteoblast differentiation [29]. Immunofluorescence staining were performed to this phenomenon. As shown in **Fig. 2i**, CD163^+^ ACP5^+^macrophage and CD163^+^ GPNMB^+^ macrophage was almost only expressed in macrophages from Randall’s plaque. Finally, we utilized Cellchat (see method) to identify unique outgoing and incoming communication of macrophage derived Randall’s plaque. As shown in **sFig. 2b and sFig. 2c**. ACKR1 and TGFBR2 is related to chemotaxis and differentiation. The GRN-SORT1 May promote mineralization of the extracellular matrix during osteogenic differentiation. Notably, SPP3 secreted by macrophage derived Randall’s plaque interact receptors expressed on T cells, mast cells and macrophage.

### COL15A1^+^ PCDH17^+^ ECs modify abnormal lipid metabolism

In total, 2783 ECs were identified in Randall’s plaque and normal renal papillae tissue and grouped into five clusters shown in **Fig. 3a, b**. Further attempts were made to determine markers of each cluster and categorize them to known ECs types. Clusters 0, 1, 2, 4 have high expression of blood endothelial cells markers (CD34, FLT1). Cluster 3 alone has high expression of lymphatic endothelial cell markers (PDPN, PROX1) (**Fig. 3c**). Cluster 3 was enriched in normal renal papillae, while cluster 0 was enriched in Randall’s plaque tissue (**Fig. 3d**). To identify Randall’s plaque-specific marker genes in ECs, genes that were elevated in Randall’s plaque-derived ECs were overlapped with those primarily expressed in ECs. Five genes, including COL15A1 and PCDH17 showed best specificity (**Fig. 3e, f**). Strikingly, Pseudotime analysis of the ECs using Monocle suggested a progression is highly correlated with decrease of COL15A1^+^ PCDH17^+^ EC (Double Positive Endothelia Cell, DPEC) percentage in the clusters (z = −2.96, p = 0.003; **Fig. 3g**). Adipogenesis pathway activation was observed in the COL15A1^+^ PCDH17^+^ subsets (**sFig. 3a**). The Cellchat tool was employed to assess the cell-cell interactions. Of note, DPECs harbored many interactions from ECs and the unique outgoing communication pairs from each cell type (**Fig. 3h, i**). Notably, DPECs contained many inferred interactions with epithelial cells. Of the DPECs unique pairs, there were interactions between SPP1, LGALS9 and CD44, which promotes cell–cell interactions, cell adhesion and crystal adhesion (**Fig. 3j, sFig. 3b**) [30, 31]. The differential gene expression analysis between DPECs and ECs shown that SPP1 is one of significantly high express genes (**sFig. 3c**). Pathway activation comparison used to further explore the function of DPECs. The activation of enriched pathways in DPECs are related to SPP1 expression (**Fig. 5k**). Together, these data suggest that DPECs modify the adipogenesis and upregulating proliferation while further induce crystal adhesion.

**Fig. 3.**
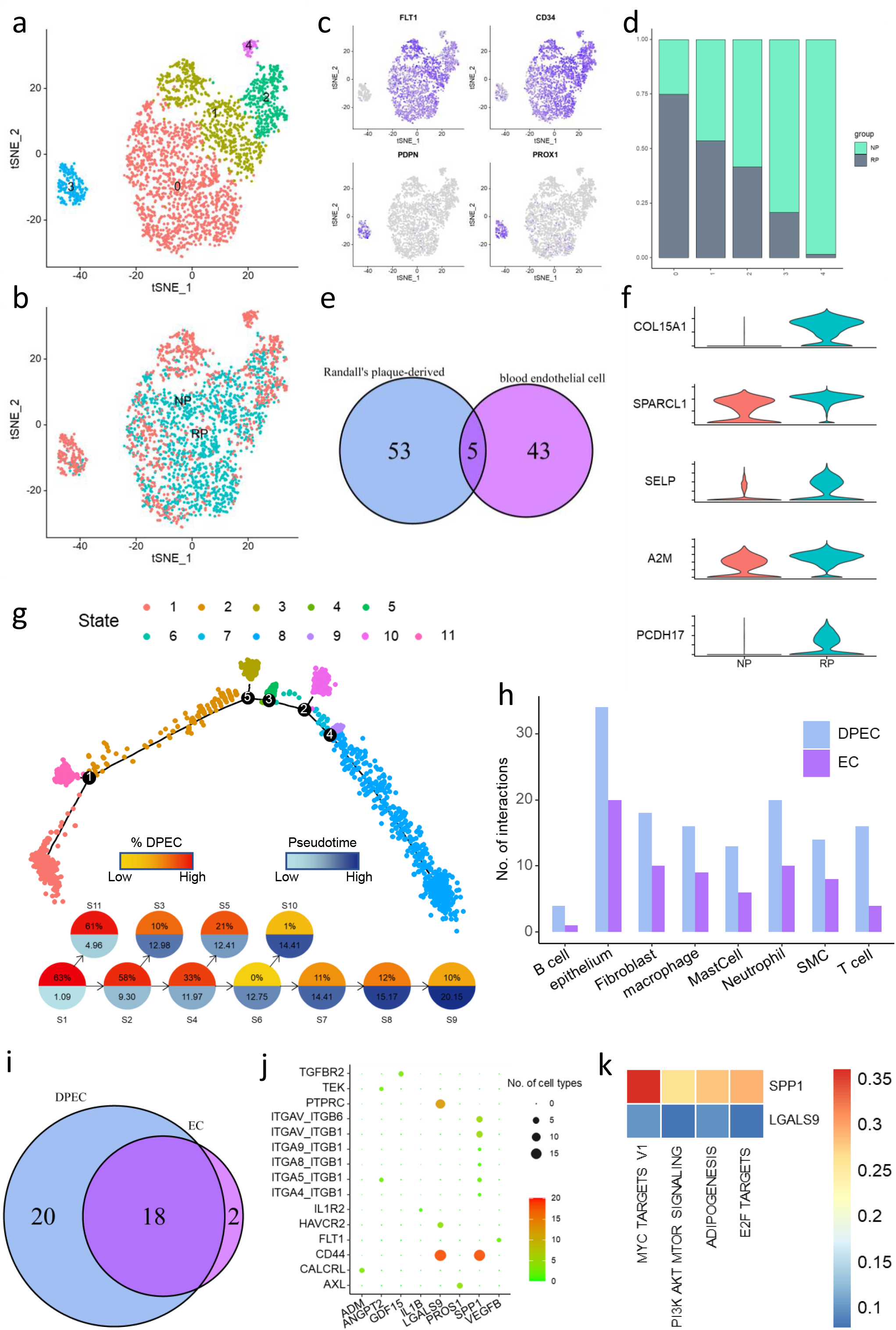
Identifying a double positive EC cell subset in ECs. a. tSNE plot of endothelial cells, color coded for 5 clusters of endothelial cells.
b. tSNE plot of endothelial cells, color coded for endothelial cells from Randall’s plaque and normal renal papillae.
c. tSNE plot of blood endothelial cell (FLT1, CD34) and lymphatic endothelial cell markers (PDPN, PROX1), while color represents expression level.
d. For five subclusters identified in ECs, the fraction of subclusters that originated from the Randall’s plaque tissue and normal renal papillae.
e. The Venn diagrams show the number of differentially and commonly markers identified between Randall’s plaque-derived ECs and all ECs.
f. Violin plots of marker genes for Randall’s plaque-derived ECs.
g. Top: all ECs from Randall’s plaque and normal renal papillae tissues ordered along pseudotime trajectories, color-coded by cluster. Bottom: schematics of the trajectories, with pie chart color-coded by average pseudotime and percentage of cells from Randall’s plaque samples in the indicated cluster. Two-sided P values are calculated for Spearman’s rank correlation and adjusted for multiple comparisons.
h. Number of unique cell communication pairs in each cell type.
i. Overlap among the unique outgoing cell communication pairs from ECs and DPECs.
j. Unique Ligand-receptor interaction from DPECs detected by Cellchat. color and size coded number of cell types.
k. heatmap to show the correlation between hallmark pathway activities and SPP1, LGALS9. color coded the Spearman’s rank correlation.

### Loop of Henle as the main sender of SPP1 signaling

We detected 7612 epithelial cells. Reclustering these cells revealed 16 clusters (**Fig. 4a, b**). Attempted were made to clarify genes specific to each of clusters and align them with known ECs types (**Fig. 4c, sFig. 4a**). This revealed 3 sets of 1799 basal cells found in Randall’s plaque and normal kidney papillae (cluster 3, 5 and 7; marker genes KRT5, TP63), and 5 sets of 2988 superficial cells (cluster 0, 1, 2, 12 and 14; marker genes UPK1B, UPK1A), and 3 sets of 1353 pelvic epitheliums (cluster 8, 9, 10, 11; marker genes SAA2, KRT23), and 2 sets of 885 loop of Henle cells (cluster 4, 13; marker genes SLC12A1, CLDN16), and 2 sets of 587 collecting duct cells (cluster 6, 15; marker genes AQP2, SLC14A2). Of these, Pelvic epithelium was particularly Randall’s plaque enriched (**Fig. 4d**). Cellchat were used to identify SPP1 signaling roles of cell groups as well as the major contributing signaling. This is consistent with previous analysis that RD macrophages, DPECs, collecting duct cells and Loop of Henle cells as sender of SPP1 signaling. Interestingly, Loop of Henle cells as the most predominant sender (**Fig. 4e**). A direct comparison of Randall’s plaque and normal papillae-derived Loop of Henle cells revealed 136 elevated and 160 genes decreased genes (adjust p value < 0.05, | log2(Fold change) | > 1) (**Fig. 4f**; **Supplement Table 2**). Notably, the top DEGs, elevated or decreased, were noisily linked to the Randall’s plaque formation. Earlier studies indicated that up-regulated LCN2 and down-regulated UMOD, SLC12A1 is essential for apatite deposits in the kidney papillae. CA12, LTF, SERPINA1 involved in lipid metabolism [8, 32]. Not only that, the expression of SPP1 was greatly improved in Randall’s plaque, compared with the renal papillae. As shown in **Fig. 4g**, the GSEA revealed that the fatty acid metabolism, adipogenesis and inflammatory response pathway were enriched in Loop of Henle cells with high expression of SPP1. This suggest lipid metabolism is the potential causes of elevated SPP1 signaling.

**Fig. 4.**
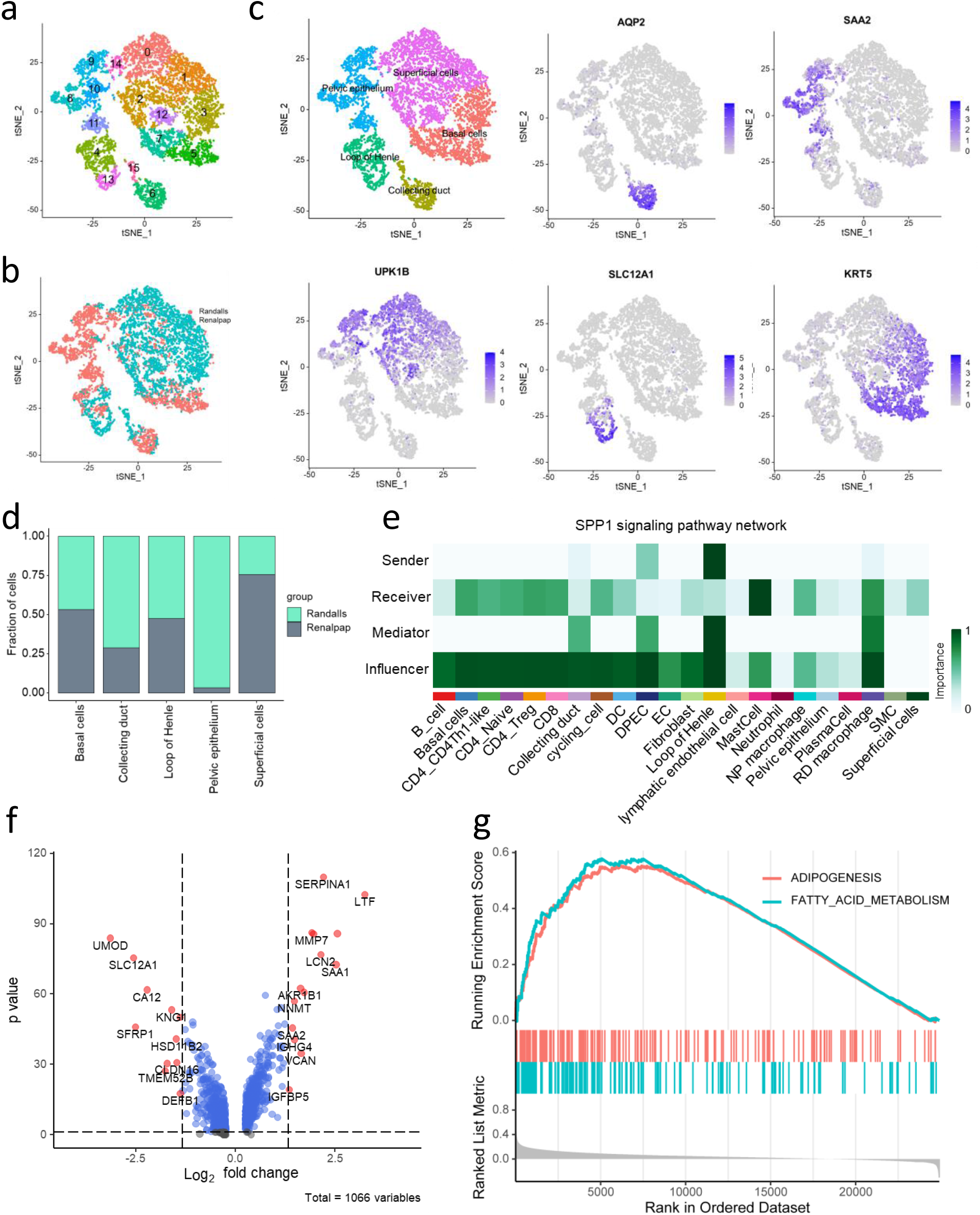
Loop of Henle cells as the most predominant sender of SPP1 signaling. a. tSNE plot of fibroblast cells, color coded for 16 subclusters of epithelial cells. Color coded by cluster.
b. tSNE plot of fibroblast cells, color coded for epithelial cells from Randall’s plaque and normal renal papillae. Color coded by group.
c. tSNE view of marker gene expression.
d. For five cell types identified in fibroblast cell, the fraction of subclusters that originated from the Randall’s plaque tissue and normal renal papillae.
e. Heatmap showing dominant senders, receivers, mediators and influencers in the intercellular communication network base on SPP1 signaling. color coded importance.
f. The Volcano plot showing differential gene expression of renal papillae Loop of Henle derived-Randall’s plaque vs derived-normal.
g. GSEA reveals two pathways enriched in SPP1 high expression of Loop of Henle cells. FDR <0.01 was considered as significantly enriched.

### Loss of universal fibroblast gene in Randall’s plaque tissue

Fibroblasts have been proven to have the potential for topic calcification in the formation of RP. But the mechanism of differentiation has not received substantial investigation. This is partly because fibroblasts show diverse phenotypes that vary with culture conditions. Herein, 2815 fibroblasts were found. Subclustering identified 5 subtypes (**Fig. 5a**). While fibroblasts were overall only modestly enriched in Randall’s plaque. cluster 0 and cluster 2 were enriched in Randall’s plaque were strongly enriched in normal renal papillae. cluster 1, 3 and 4 were found almost exclusively in normal renal papillae (**Fig. 4b, c**). Each fibroblast expresses distinct extracellular matrix biomolecules and collagens, example. e.g., cluster 1 express COLCA1 and cluster 2 express COL3A1 (**Fig. 5d**). In contrast to normal renal papillae, Randall’s plaque expresses high COL3A1, COL6A3 (collagens type III, VI), low levels of PCOLCE2, which are associated with trimerization of collagen chain and degradation of the extracellular matrix. Each collagen has a distinct function, suggesting that fibroblast clusters have developed functional specialization [33].

RGS5, as myo-associated fibroblast maker, showed high expression in cluster 0, 2. PDGFRA^+^ fibroblasts express high levels of chemokines and cytokines, which is similar to inflammatory fibroblasts (**Fig. 5f**) [34, 35]. To identify specific markers for NP specific fibroblast (cluster 1, 2, 4), genes that were primarily expressed in fibroblast cells were overlapped with those elevated in NP specific fibroblast cells. A total of 9 genes were identified, among which OGN and C7 showed the best specificity (**Fig. 5g**). We noticed high expression of MGP in NP specific fibroblast, which is a blocker of many bone morphogenetic proteins (BMPs) [36, 37].

Results from the GSVA analysis showed that the Inflammation-related, Proliferation-related and stress-related signaling pathways were enriched in RGS5^+^ fibroblasts, especially in cluster 2, which mostly derived from Randall’s plaque tissue. SCENIC revealed that genes regulated by the JUN, JUND, EGR1, FOSB and ATF3 transcription factors were upregulated in RGS5^+^ fibroblasts. Notably, ATF3 is involved in the complex process of cellular stress response, while FOSB and EGR1 as regulators of cell proliferation, differentiation, and transformation. Fos/Jun can trigger inflammation responses in fibroblasts (**Fig. 5h**). Immunofluorescence double staining images and the R value of overlap coefficient revealed the expression of OGN and C7 were markedly increased in NP-fibroblast compared with RP-fibroblast (**Fig. 5i, sFig. 5**). These data reveal NP specific fibroblast (COL1A1^+^ C7^+^ OGN^+^ fibroblast) suppresses amorphous CaP transforms into hydroxyapatite and carbonate apatite. Matthew B. Buechler et al suggested activation in human fibroblasts were associated with loss of universal fibroblast gene expression [38]. The absent of COL1A1^+^ C7^+^ OGN^+^ fibroblast is potential causes of Randall’s plaque.

**Fig. 5.**
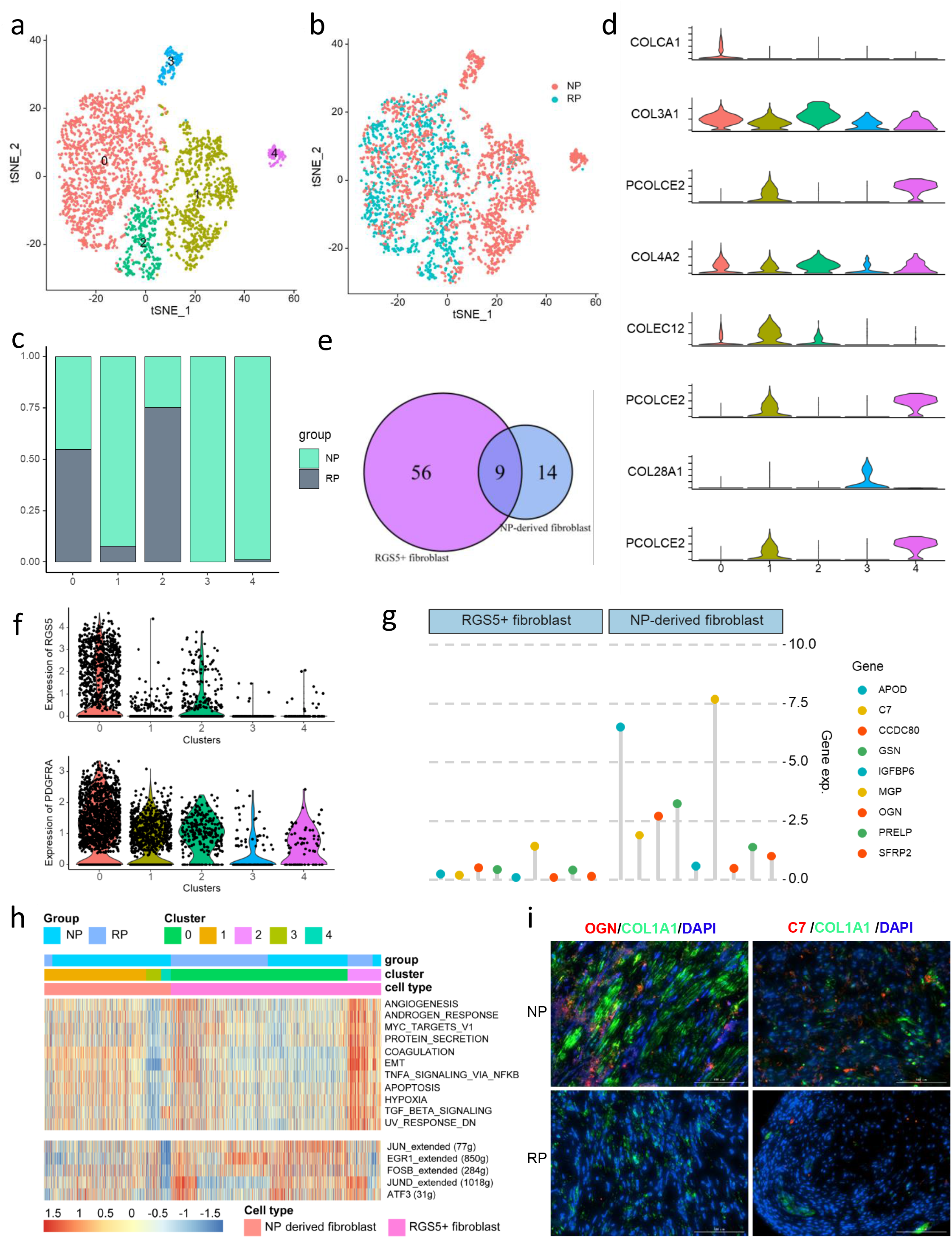
fibroblast in Randall’s plaque lack suppressing amorphous CaP transforms into hydroxyapatite potential. a. tSNE plot of fibroblast cells, color coded for 5 subclusters of fibroblast cells.
b. tSNE plot of fibroblast cells, color coded for fibroblast cells from Randall’s plaque and normal renal papillae.
c. For five subclusters identified in fibroblasts, the fraction of subclusters that originated from the Randall’s plaque tissue and normal renal papillae.
d. Violin plots of collagens and extracellular matrix molecules.
e. The Venn diagrams show the number of differentially and commonly markers identified between Randall’s RGS5+ fibroblast and NE-derived fibroblast.
f. Violin plots of fibroblast-associated makers for five subgroups.
g. Relative expression of 9 common differential genes.
h. Top: Differences in 50 hallmark pathway activities scored with GSVA. Shown are t values calculated by a linear model. Bottom: Heatmap of the area under the curve (AUC) scores of expression regulation by transcription factors estimated by SCENIC.
i. Immunofluorescence double staining with antibodies against OGN and BMSP (red immunofluorescence) with anti-COL1A1 (green immunofluorescence) in the same slide of Randall’s plaque or normal renal papillae. (DAIP: blue immunofluorescence)

### Constructing an SPP1-based regulatory network for Randall’s plaque

Further, cell–cell interaction networks were studied using the Cellchat. RP macrophages, loop of Henle and DPECs exhibited the highest number of interactions with other cell types, especially with RGS5^+^ fibroblasts (**Fig. 6a**). Based on the expression abundance, and results from SCENIC and GSVA a, data concerning the interaction pairs such as SPP1, complement, VEGF, EGF, PTN and CXCL families were obtained. RD macrophages, loop of Henle as the main source of SPP1 signaling. The receptors of which includes CD44, ITGAV, ITGB1, ITGB5, ITGB6, ITGA4, ITGA5, ITGA8. Notably, RD macrophage and DPECs also receive signals from SPP1 secreted by DPECs. The CXCL2/3/8-ACKR1 signature was more intense in the DPECs compared with ECs, specially from RD macrophage. DEPCs exhibit elevated CXCL12 expression, and its receptors, CXCR3 and CXCR4. Moreover, CXCL12 secreted by DPECs triggers immune cell infiltration in Randall’s plaque. (**Fig. 6b**). These cytokines have been linked to Randall’s plaque formation, and we pinpointed their origin. We also suggested that epithelial cell, fibroblast and RD macrophages secrete VEGF (VEGFA and VEGFB), which binds to VEGFR1 and VEGFR2 on DPECs to promote angiogenesis. SDCs were expressed on Loop of Henle and RD macrophage and EGF family were expressed on Loop and Henle. These receptors could bind to PTN and show pro-proliferation effects. Notably, the ITGAX and ITGB1 on RD macrophage receives C3 signals from epithelial cells. Previous study showed C3 has both pro-inflammatory and adhesion effects in Randall’s plaque. (**Fig. 6c**). To summarize the above, Loop of Henle is major source of SPP1 signaling. DEPCs and RD macrophage acts as a core of the SPP1 signaling pathway, which induce spontaneous interstitial CaP crystal and tissue inflammatory formation. (**Fig. 6d**).

**Fig. 6.**
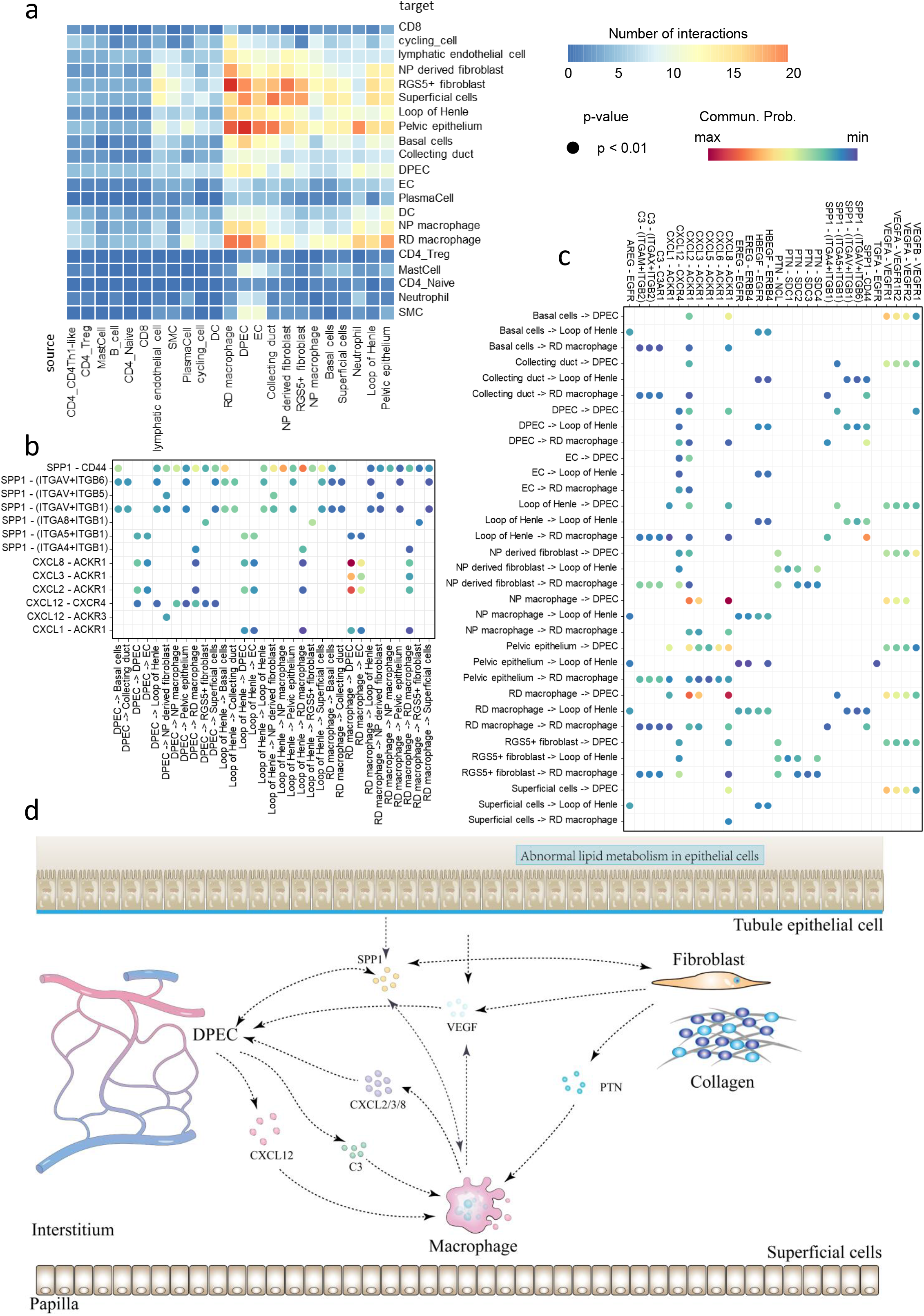
Cell-cell communication network in Randall’s plaque tissue micro-environment. a. Heatmap show number of potential ligand-receptor pairs between cell groups predicted by Cellchat.
b. Bubble plots show outgoing communications patterns of DEPCs, Loop of Henle and RP macrophage.
c. Bubble plots show incoming communications patterns of DEPCs, Loop of Henle and RP macrophage.
d. Predicted regulatory network base on DEPCs, Loop of Henle and RP macrophage.

## Discussion

Increasing evidence has shown that Randall’s plaque is caused by interactions among the external environmental factors and internal demic factors. Recent studies have begun to uncover that inflammation and oxidative stress lead to the RP initiation. Even some researchers have focused on how macrophage polarization and recruitment influence RP formation. However, only little is known about the characteristics of tissue microenvironment in Randall’s plaque. This necessitates further investigations into the changes of cell type and cell communication. Thus, in the current work, we investigated the differences in tissue micro-environment between Randall’s plaque and normal renal papillae, showed the complete list of stromal cells, epithelial cells and immune cells in Randall’s plaque tissue and clarified markers linked to each cell subgroup at the protein level. Herein, we revealed Randall’s plaque-associated cell subgroups, pathway, and networks. These results will inform our understanding novel cell subgroups in microenvironment linked to Randall’s plaque and provide a new perspective for further studies on Randall’s plaque.

Macrophages drive the formation of interstitial crystals in Randall’s plaque have long been known. Such macrophages have various phenotypes which are influenced by the disease stage, type and microenvironment [39]. Previous studies have identified that genes associated with inflammatory macrophage were upregulation and genes associated with pro-inflammatory macrophage were downregulation [23, 24]. However, the results of their study were not consistent with ours. We used two different bioinformatics approach to assess degree and direction of macrophage differentiation. As shown in our results, a visible increase in the number of infiltrating M1 and M2-like macrophage was presented in both Randall’s plaque and normal renal papillae. However, there is no evidence that the imbalance in differentiation between M1 and M2. An image-base immunofluorescence was used to validate this result. The phenomenon that the expression of osteoblast-associated genes increased markedly suggest that osteogenic microenvironment was created in Randall’s plaque. In addition, we have noted a key observation. The GPNMB+CD163+ and ACP5+ CD163+ macrophage almost exclusively observed in Randall’s plaque. Thus, the conventional marker of macrophage was not efficient in identifying macrophage subgroups. anti-inflammatory effects of GPNMB have also been observed in macrophage. ACP5 is one of the genes reported to be associated with bone metabolism. These results suggest that macrophage in Randall’s plaque have additional regulation function of bone metabolism.

Second, we identified COL15A1 and PCDH17 as makers that distinguishes activated endothelial from other cell types in renal papillae, including endothelial from normal renal papillae [40]. COL15A1 is involved in fibrosis development in cases of hepatocellular and biliary damage. Portal myofibroblasts improve remodeling of the vasculature leading to the genesis of cirrhosis driven by the secreted microparticles. PCDH17 code potential calcium-dependent cell-adhesion protein. DPECs exhibit abundance of cell-cell communications that is not available in most normal endothelial. these special cell to cell communications involved in osteopontin, collagens, and matrix metalloproteinases. By pseudotime analysis, revealed Randall’s plaque formation was accompanied by differentiation of endothelial cells. these results suggesting that alterations in osteogenicity in the endothelial cells is one of most important cause of contribute to RP formation. The first differentiation path for impaired ECs function in RP was examined and pathways involved in various stages of its development. Inhibiting this pathway could suppress ECs impairment. This could be an effective approach to block formation of Randall’s plaque. Moreover, a novel TF was uncovered which may expand our understanding of its pathological mechanisms.

Thirdly, the gene (SPP1) that code for osteopontin (OPN) is considered to play interact with other genes involved in lithogenesis to regulate stone matrix. A recent study suggest that RPs are accompanied by apoptosis tubular/ interstitial/urothelial/ as well as OPN aggregation, as a result of oxidative stress and inflammation [8, 41]. But results of the studies in animal models indicated that osteopontin prevents CaOx crystallization in kidney [42]. Our results show that SPP1 is overexpressed on multiple cell types in Randall’s plaque compare with normal renal papillae. cell-cell communication revealed that SPP1 interact with CD44 which is crystal-related genes. In fact, SPP1 pathway may be crucial signaling pathway the process of Randall’s plaque formation. Not only that, we also found the Loop of Henle, DPECs, collect duct and RD macrophages is main sender of SPP1 signaling. The above findings may contribute to improve our understanding of the role of SPP1 signaling in Randall’s plaque microenvironment.

In addition, dyslipidaemia is liked to kidney stone formation [43]. previous studies showed obese model mice and rats have significantly larger amounts of renal crystal deposition compared with lean controls. In mice, FABP4 knockout triggered urinary and renal crystals. In this study, abnormal lipid accumulation was observed in vascular endothelial cells and Loop of Henle. This might be an explanation for uric acid and CaOx stone formers remove a higher amount of lipids compared to individuals without stones [44].

The following limitations need to be recognized study. Firstly, as limitations in sc-seq sample size, the findings presented here may not apply to all type of Randall’s plaque microenvironment. This study used sequencing samples from patients with calcium oxalate stones. tissue microenvironment from patients who had other composition stone may not be the same. Second, the data may be affected by sample collection bias. Although we confirm sc-seq sample obtained from a plaque area. tissue depth could not be accurately controlled due to manual manipulation. Third, preparation of single cell in suspension required the digestion for cell. part of cells was filtered out. This may result in some information loss. In addition, suitable control samples should have comprised normal renal papillae tissue. But we unable to obtain this tissue from healthy volunteer due to ethical restrictions. here, we have used normal renal papillae from localized renal carcinoma patients as controls.

Collectively, the transcriptome landscape of Randall’s plaque was uncovered in this study. Further larger single-cell sequencing of Randall’s plaque is needed to identify potential tissue factors of stone recurrence.

## Supporting information

Supplement figure 1, Phenotypic characterization of macrophage sub-clusters.

Supplement figure 2, Specific cell communication of RP macrophage.

Supplement figure 3, Specific cell communication of DPECs.

Supplement figure 4, tSNE plot of endothelial cell markers. Color coded expression of marker.

Supplement figure5. The expression of OGN and BMSP were markedly increased in NP-fibroblast compared with RP-fibroblast.

Supplementary table 1: Cell characteristics in renal papillae.

## Declarations

Ethics approval and consent to participate

The patient data in this work were acquired from the publicly available datasets whose informed consent of patients were complete.

## Consent for publication

Not applicable.

---

**Supplement figure 1, Phenotypic characterization of macrophage sub-clusters.**

a. tSNE plot of macrophage, color coded for relative expression of macrophage markers.
b. Percent of two categories macrophage between Randall’s plaque and normal renal papillae tissue.
c. Left, Violin plots of M1 signature between NP and RP. Middle, Violin plots of M2 signature between NP and RP. Right, Ratio of M1 and M2 signature between NP and RP.
d. Violin plots of bone metabolism activity among five clusters.

**Supplement figure 2, Specific cell communication of RP macrophage.**

a. Violin plots of cell markers between NP and RP.
b. Bubble plot of specific incoming cell communications of RP macrophage. Color and size coded possibilities.
c. Bubble plot of specific outgoing cell communications of RP macrophage. Color and size coded possibilities.

**Supplement figure 3, Specific cell communication of DPECs.**

a. Bubble plot of relationship between adipogenesis activation and pseudotime. Size and color coded the number of cells.
b. The Volcano plot showing differential gene expression of DEPCs vs ECs.
c. Circle plot of specific cell communications of EPECs by counting the number of links.

**Supplement figure 4,** tSNE plot of endothelial cell markers. Color coded expression of marker.

**Supplement figure5. The expression of OGN and BMSP were markedly increased in NP-fibroblast compared with RP-fibroblast.** a, Top, Immunofluorescence double staining of OGN and COL1A1 between normal renal papillae and Randall’s plaque tissue slide. Bottom, Immunofluorescence double staining of PSMB and COL1A1 between normal renal papillae and Randall’s plaque tissue slide. Image pro plus used to calculate Pearson correlation coefficient and overlap coefficient. b, Violin and box plots for the distribution of overlap coefficient.

**Supplementary table 1:** Cell characteristics in renal papillae.

**Supplementary table 2: Differential gene expression between Randall’s plaque and normal renal papillae tissue.**

