## Supplementary figures and images for "Landscape of microenvironment in Randall’s plaque by single-cell sequencing"

### Supplement figure5. The expression of OGN and BMSP were markedly increased in NP-fibroblast compared with RP-fibroblast.

#### Supplement figure 5

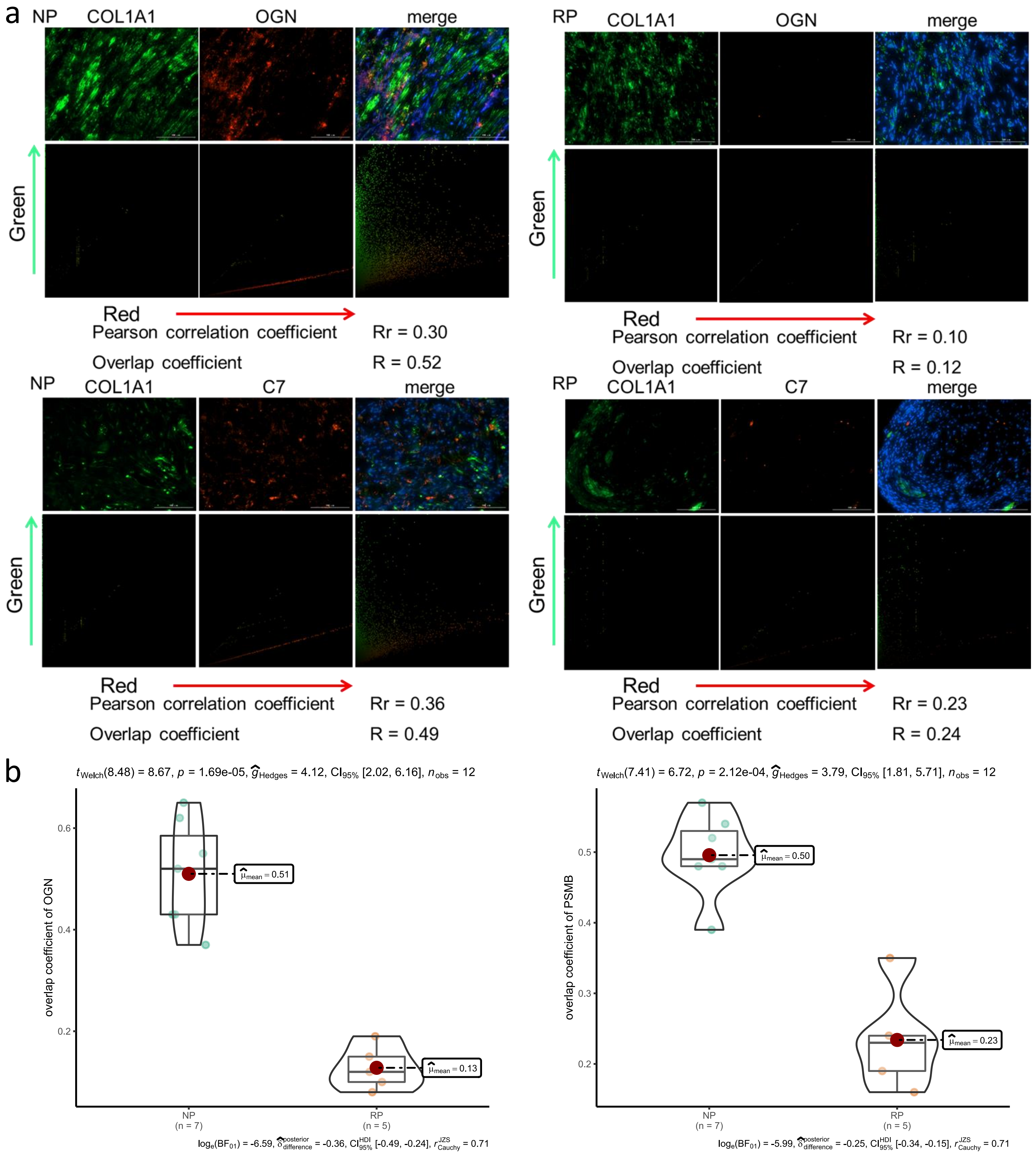

### Supplement figure 1, Phenotypic characterization of macrophage sub-clusters.

#### Supplement figure 1

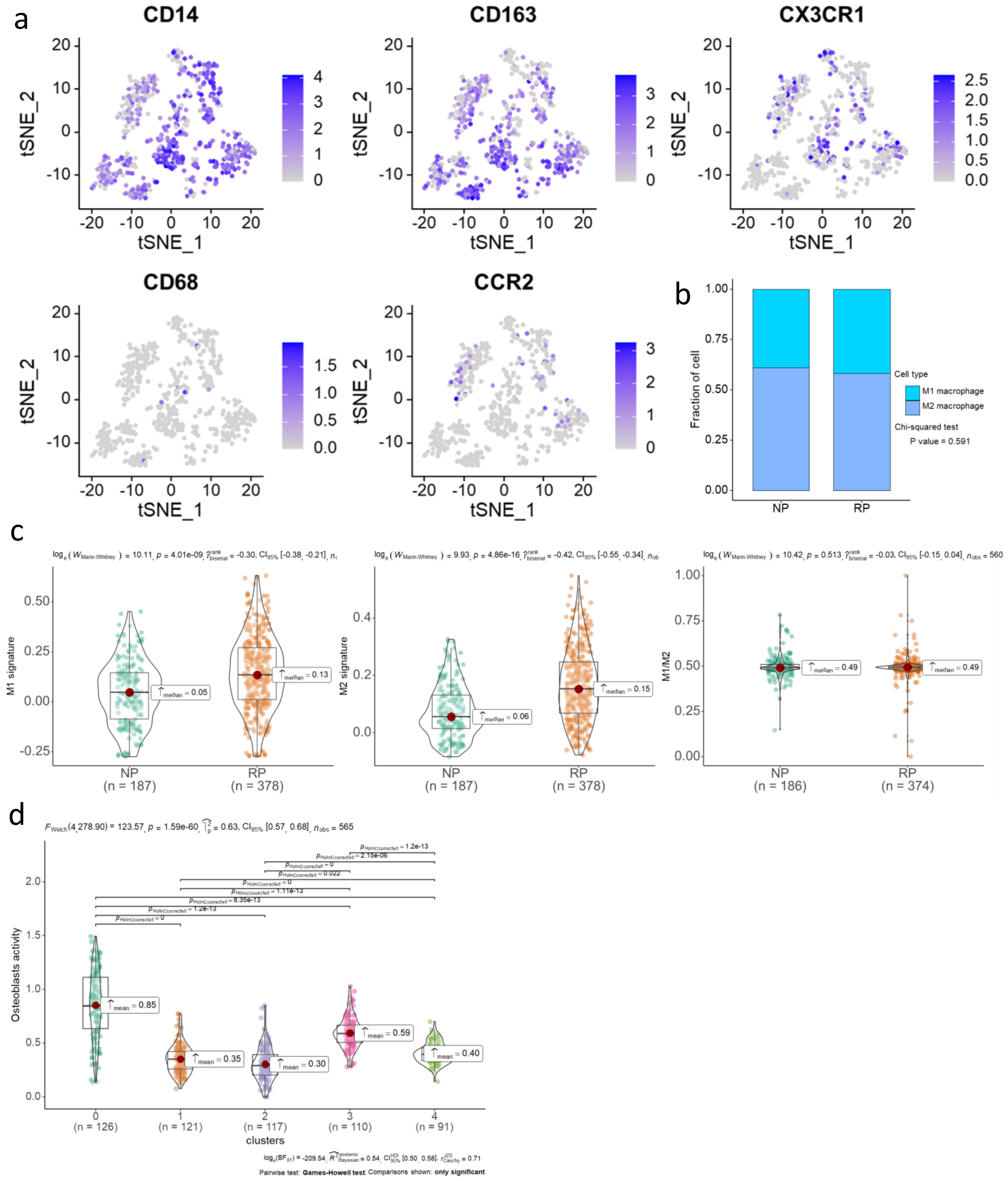

### Supplement figure 2, Specific cell communication of RP macrophage.

#### Supplement figure 2

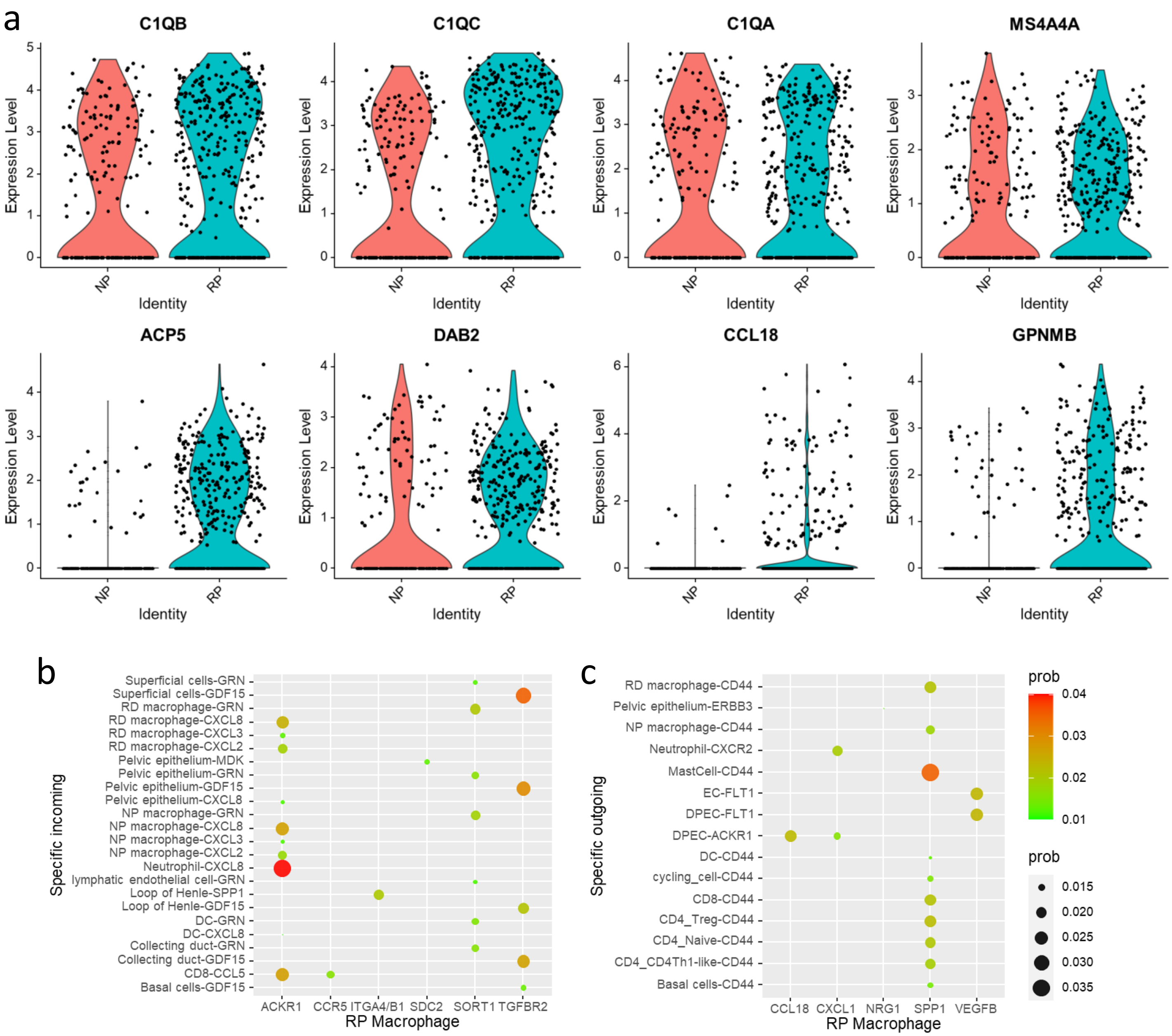

### Supplement figure 3, Specific cell communication of DPECs.

#### Supplementary figure 3

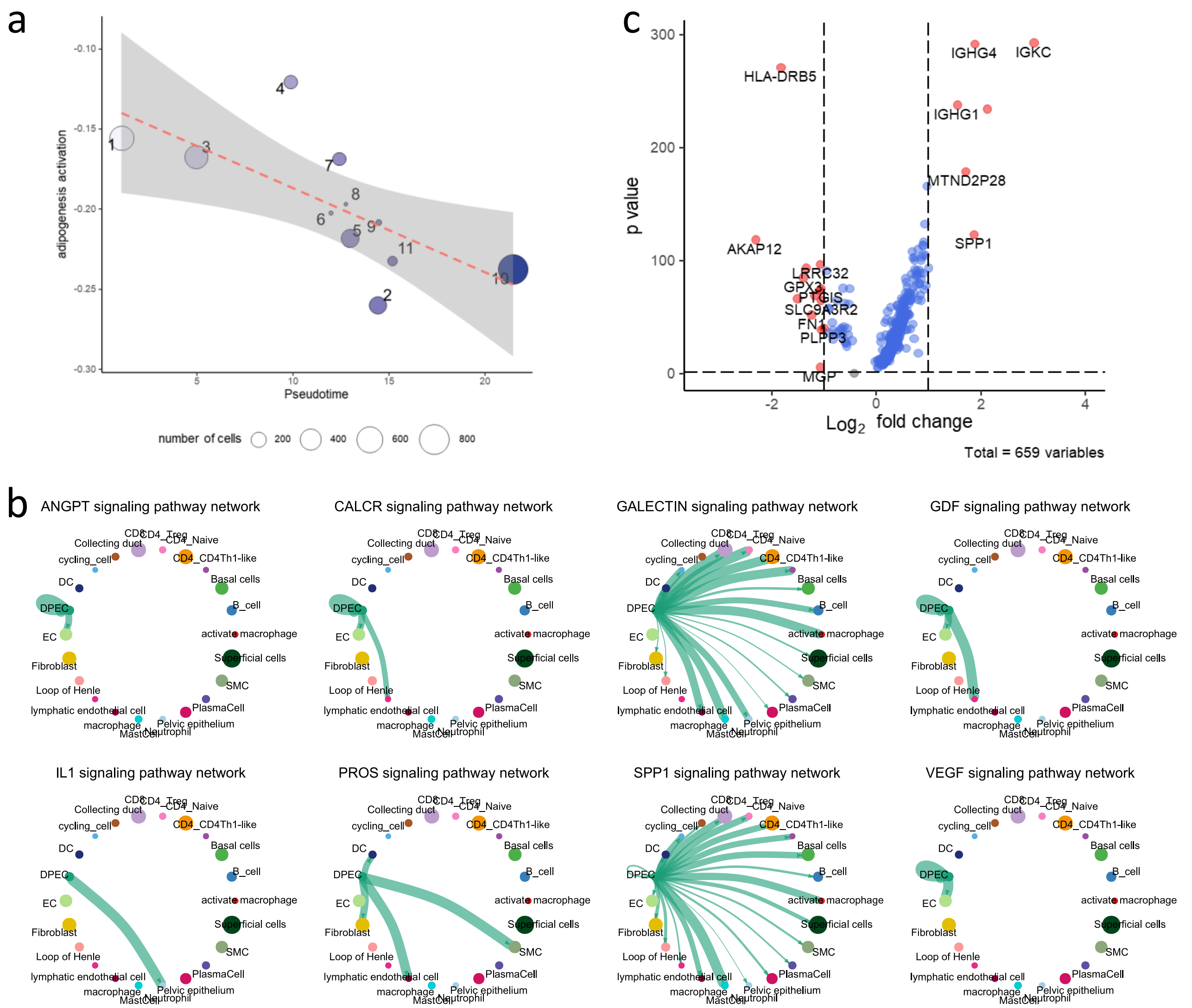

### Supplement figure 4, tSNE plot of endothelial cell markers. Color coded expression of marker.

#### Supplement figure 4

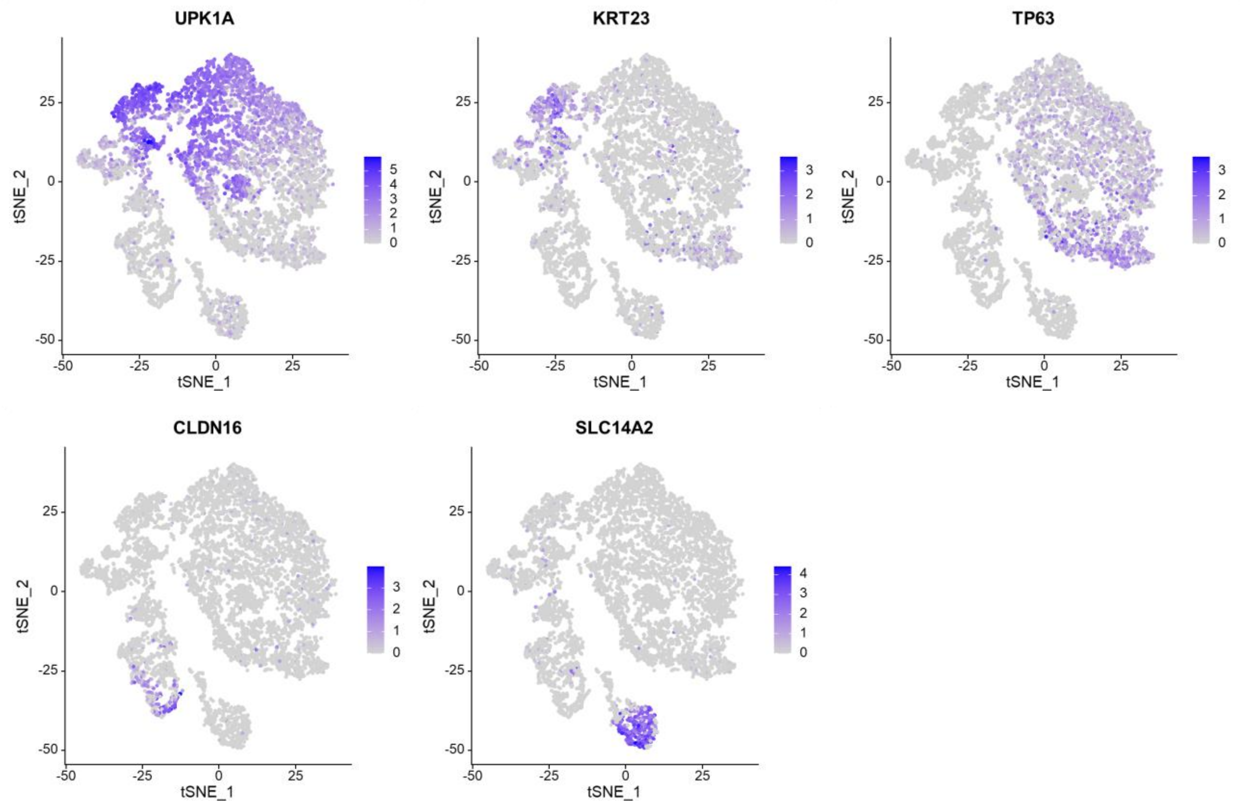
