## Supplementary table 1: Cell characteristics in renal papillae. for "Landscape of microenvironment in Randall’s plaque by single-cell sequencing"

| CELL TYPE | CELLS | GENES | UMIS | KNOWN MARKERS | POTENTIAL MARKER GENES |
|---|---|---|---|---|---|
| **B cell** | 1106 | 937.6347 | 1944.785 | CD19 | CD83, MS4A1, CD79A |
| **Basal cells** | 1799 | 2388.291 | 7045.721 | KRT5, PVRL4, TP63 | SLC2A1, SFN, AQP3, KRT5, KRT17, S100A2, CLDN1, MIR205HG, TRIM29, ERRFI1, SOWAHC |
| **CD4_CD4Th1-like** | 166 | 1099.518 | 2088.337 | STAT3, AHR, BATF | RGS1, SPOCK2, IL7R, PRDM1 |
| **CD4_Naive** | 2231 | 972.4043 | 2115.016 | ABLIM1, CCR7 | |
| **CD4_Treg** | 437 | 1024.611 | 1888.721 | FOXP3, CTLA4 | TIGIT, CTLA4, ICOS |
| **CD8** | 2217 | 940.6297 | 1795.126 | | CCL5, CD8A, CCL4, GZMA, CST7 |
| **Collecting duct** | 587 | 2636.169 | 7336.041 | AQP2, SLC14A2 | DEFB1, MAL, AIF1L, ATP1B1, KCNJ16, EPCAM, SCNN1A, MUC1, AQP2, HSD11B2, CDH16, SLC14A2, AOC1, FXYD4, AKR1B1, TMEM213, L1CAM, STC1, ST6GAL1, PAX8, TSPAN1, SCIN, CLDN8, RDH10, FOLR1, SLC5A3, SLC16A5, TFCP2L1, MYH10, TMEM176B, TMEM176A, AC023421.1, MAL2, ADGRF1, CD82 |
| **Cycling cell** | 76 | 2867.618 | 9520.105 | | MKI67, TOP2A, CENPF, NUSAP1, TYMS, TPX2, RRM2, ASPM, BIRC5, CENPE, STMN1, HIST1H4C, HMGB2 |
| **DC** | 650 | 1551.822 | 4469.06 | CD1C, FCER1A | HLA-DMB, HLA-DPB1, GPR183, HLA-DQA1, HLA-DQB1, C1orf162 |
| **EC** | 2585 | 1835.019 | 4746.61 | CD34, FLT1 | PLVAP, ACKR1, FLT1, CD34, PECAM1, AQP1, EMCN, SPARCL1, TM4SF1, RNASE1, CD93, VWF, ADGRL4, TSPAN7, MMRN2, CALCRL, IFI27, SLCO2A1, PTPRB, PALMD, CLEC14A, HEG1, ADGRF5, SOX18, ESAM, CDH5, JAM2, KDR, LDB2, TM4SF18, ENG, CYYR1, FAM198B, PCAT19, A2M, ERG |
| **Fibroblast** | 2003 | 2050.214 | 5117.761 | COL1A1, COL3A1, RGS5, PDGFRA | SPARC, NR2F1, COL14A1, DCN, COL1A2, COL1A1, LUM, FBLN1, C1S, CCDC80, COL3A1, MMP2, PTGDS, C1R, PRELP, SERPINF1, PDGFRA, ISLR, CRISPLD2, CTSK, COL6A2, COL6A3, VCAN, COL6A1, SRPX, APCDD1, OLFML3, TIMP2, MFAP4, PCOLCE, COL8A1, LRP1, DDR2, FSTL1, LTBP1, ANTXR1, OSR2, ITGA8, COL5A1, CYBRD1, CDH11, TGM2, FBLN5, MATN2, AEBP1, FOXF1, CYGB |
| **Loop of Henle** | 885 | 2723.698 | 8522.028 | SLC12A1, CLDN16 | SPP1, CD24, CXCL14, CRYAB, POU3F3, CLDN10 |
| **lymphatic endothelial cell** | 198 | 1659.864 | 4139.025 | PDPN, PROX1 | RAMP2, LMO2, TIE1, GNG11, CAVIN2, MMRN1, CCL21, PROX1, FABP4, FLT4, COLEC12, RELN, TFF3, MRC1, PKHD1L1, CHRDL1, NRP2, TFPI, FLRT2, LAMA4, MYCT1, PDPN, EFEMP1, AKAP12, ABI3BP, |
