## Supplemental Data 1 for "Landscape of microenvironment in Randall’s plaque by single-cell sequencing"

**Supplementary Table 2**: Differential gene expression between Randall's plaque and normal renal papillae tissue.

| Gene | p_val | avg_logFC | pct.RP | pct.NP | p_val_adj |
|---|---|---|---|---|---|
| LTF | 2.87E-103 | 3.177819 | 0.836 | 0.095 | 1.09E-98 |
| IGKC | 1.60E-86 | 2.399898 | 0.879 | 0.339 | 6.10E-82 |
| SAA1 | 2.19E-73 | 2.33386 | 0.583 | 0.009 | 8.33E-69 |
| SERPINA1 | 8.56E-111 | 2.148196 | 0.95 | 0.289 | 3.26E-106 |
| LCN2 | 1.31E-77 | 1.974912 | 0.667 | 0.059 | 4.97E-73 |
| SLPI | 5.41E-86 | 1.897579 | 0.875 | 0.209 | 2.06E-81 |
| MMP7 | 1.17E-86 | 1.869084 | 0.944 | 0.367 | 4.43E-82 |
| C3 | 1.83E-61 | 1.588955 | 0.641 | 0.128 | 6.95E-57 |
| AKR1B1 | 3.83E-63 | 1.562148 | 0.823 | 0.393 | 1.46E-58 |
| VCAN | 2.22E-35 | 1.502127 | 0.527 | 0.168 | 8.44E-31 |
| NNMT | 1.59E-57 | 1.342588 | 0.654 | 0.154 | 6.04E-53 |
| HSD11B2 | 1.32E-41 | -1.34148 | 0.117 | 0.514 | 5.01E-37 |
| KNG1 | 4.68E-54 | -1.49921 | 0.058 | 0.54 | 1.78E-49 |
| TMEM52B | 5.08E-28 | -1.57228 | 0.009 | 0.251 | 1.93E-23 |
| S100A2 | 3.81E-31 | -1.59877 | 0.283 | 0.588 | 1.45E-26 |
| CA12 | 8.10E-63 | -2.14265 | 0.367 | 0.763 | 3.08E-58 |
| SFRP1 | 9.82E-47 | -2.3275 | 0.078 | 0.488 | 3.73E-42 |
| SLC12A1 | 2.65E-76 | -2.48132 | 0.173 | 0.746 | 1.01E-71 |
| UMOD | 1.35E-84 | -3.07858 | 0.071 | 0.718 | 5.11E-80 |
